# A neural correlate of visual discomfort from flicker

**DOI:** 10.1101/2020.01.25.919472

**Authors:** Carlyn Patterson Gentile, Geoffrey K. Aguirre

## Abstract

The theory of “visual stress” holds that visual discomfort results from overactivation of the visual cortex. Despite general acceptance, there is a paucity of empirical data that confirm this relationship, particularly for discomfort from visual flicker. We examined the association between neural response and visual discomfort using flickering light of different temporal frequencies that separately targeted the magnocellular, parvocellular, and koniocellular post-receptoral pathways. Given prior work that has shown larger cortical responses to flickering light in people with migraine, we examined 10 headache free people and 10 migraineurs with visual aura. The stimulus was a uniform field, 50 degrees in diameter, that modulated with high-contrast flicker between 1.625 and 30 Hz. We asked subjects to rate their visual discomfort while we recorded steady state visually evoked potentials (ssVEP) from primary visual cortex. The peak temporal sensitivity ssVEP amplitude varied by post-receptoral pathway, and was consistent with the known properties of these visual channels. Notably, there was a direct, linear relationship between the amplitude of neural response to a stimulus and the degree of visual discomfort it evoked. No substantive differences between the migraine and control groups was found. These data link increased visual cortical activation with the experience of visual discomfort.

## Introduction

Some visual stimuli are uncomfortable to view. High-contrast spatial and temporal patterns (i.e., stripes and flicker) have this property (Wilkins, 1995), particularly when stimulus power is concentrated at frequencies in the mid-range of human perception (Fernandez & Wilkins, 2008). Stimuli with these mid-range temporal and spatial frequencies generally evoke larger visual cortex responses (Regan, 1983; Tyler, Apkarian, Levi, & Nakayama, 1979), and are detected more easily by human observers when presented at low contrast (Robson, 1966). These findings have led to the general proposal that visually uncomfortable stimuli are the result of “excessive” cortical activity (Aurora & Wilkinson, 2007). This account finds further support in the observation that people with migraine have both greater discomfort from flickering light (Yoshimoto et al., 2017) and an enhanced visual cortex response to these stimuli (Datta, Aguirre, Hu, Detre, & Cucchiara, 2013; Shibata, Yamane, Nishimura, Kondo, & Otuka, 2011).

Beyond this general proposal, however, the link between specific stimulus properties, visual discomfort, and cortical response is less clear. Only a weak or even absent correlation between visual discomfort and evoked cortical response in people without migraine has been found for stimuli varying in spatial frequency (Huang, Cooper, Satana, Kaufman, & Cao, 2003; O’Hare, 2017). Studies of sensitivity to patterns and temporal flicker have generally been made using black-and-white patterns, which probe only a small set of possible stimulus properties that may be associated with visual discomfort. The cortical visual system receives input via three different post-receptoral channels, each with different spectral (i.e., “color”) sensitivity. The magnocellular (“black-white”), parvocellular (“red-green”), and koniocellular (“blue-yellow”) post-receptoral pathways also have different spatial and temporal response properties (Kelly, 1974). Prior studies have found a relationship between reported discomfort and the chromatic contrast of spatial gratings (as expressed as distance in hue spaces; Haigh et al., 2013; Juricevic, Land, Wilkins, & Webster, 2010), suggesting that multiple post-receptoral channels contribute to visual discomfort. However, no study has examined the interaction of temporal flicker and post-receptoral channel in the induction of visual discomfort. Recent work has found that pulses of colored light from a dark background differ in the degree to which they evoke discomfort in people who are in the midst of a migraine headache (Noseda et al., 2017; Noseda et al., 2016). While again suggesting that the chromatic content of a stimulus influences visual discomfort, studies of this kind are unable to identify the contribution of the different post-receptoral pathways to discomfort. As opposed to a flash of light from darkness, a time-varying modulation of the spectral content of light around a photopic background may be used to selectively target the post-receptoral visual pathways using the principle of silent substitution (Estévez & Spekreijse, 1982). This approach is well suited to explore the relationship between visual discomfort and visual cortex responses.

Here we examine the association between visual discomfort and evoked response in primary visual cortex using flicker of different temporal frequencies that separately target the magno, parvo, and koniocellular post-receptoral pathways. Our goal was to test the hypothesis that greater neural response evoked by any particular stimulus would be associated with a report of greater visual discomfort. We obtained data from 20 subjects: ten with migraine with visual aura (MwA), and ten headache free controls (HAf). We did not observe a difference between the groups in the cortical response to our stimuli. We did, however, find a remarkably close relationship between the tendency of each stimulus to produce visual discomfort and the magnitude of evoked cortical response.

## Methods

This study was pre-registered on the Open Science Framework (https://osf.io/f7n3x/).

### Subjects

Subjects were ages 25-41 years old and recruited from the greater Philadelphia area and University of Pennsylvania campus, in many cases using advertising on digital social media services. All candidate subjects underwent screening using the Penn Online Evaluation of Migraine (Kaiser, Igdalova, Aguirre, & Cucchiara, 2018), which implements an automated diagnostic survey using the International Classification of Headache Disorders (ICHD)-3 criteria. We identified ten subjects who met criteria for a diagnosis of migraine with visual aura (MwA). To be included in the study, MwA subjects also needed to report ictal visual discomfort, as determined by a score of 6 or greater on the Choi visual sensitivity scale (Choi et al., 2009), and a response of ‘yes’ to the question in the Choi instrument regarding the presences of light sensitivity during headache free periods. We also studied ten control participants who were either entirely headache free, or had a history only of mild, non-migrainous headache. Control subjects were required to have no known family history of migraine, and no history of childhood motion sickness. Finally, controls subjects had to score 7 or lower on the Conlon Visual Discomfort Score (VDS) survey (Conlon, Lovegrove, Chekaluk, & Pattison, 1999). There was no required score for the MwA participants, although we found that these subjects reported higher visual discomfort as measured by this instrument. Table 1 summarizes the demographic information and survey results for the participants. The study was approved by the Institutional Review Board of the University of Pennsylvania. All subjects provided informed written consent, and all experiments adhered to the tenets of the Declaration of Helsinki.

**Table 1:** Subject demographics. The median and range are shown for age, sex, visual discomfort scale (VDS), and number of headache days in the past 3 months. P-values represent two-tailed t-test.

|  | MwA (n=10) | HAf (n=10) | p-value |
| --- | --- | --- | --- |
| Age (yrs) | 34 (25-37) | 28 (25-41) | 0.9 |
| Sex | M 1, F 9 | M 4, F 6 | 0.1 |
| VDS | 15 (5-36) | 4 (0-6) | 0.001 |
| # HA days<br>/3 months | 11 (0-30) | 1 (0-4) | 0.0008 |

The demographics of the two groups did not differ significantly by sex (p=0.2), although there were more women than men in the study, consistent with the known demographics of migraine (Goadsby, Lipton, & Ferrari, 2002). The Conlon VDS score was higher in the MwA group (p=0.001). The MwA subjects had a broad range of disease burden, with a median of 11 headache days in the past 3 months (range 0-30 headache days over 3 months), which was significantly higher than the median number of headache days in the headache free group (p=0.0008). None of the MwA subjects were on preventative migraine medications. One HAf subject reported the use of daily magnesium, and two HAf subjects reported the use of serotonin-reuptake inhibitors, presumably for management of mood.

Candidates were excluded for a history of glaucoma, generalized epilepsy, a concussion in the last 6 months, or on-going symptoms from head trauma/concussion. Participants were excluded if best corrected distance acuity was below 20/40 based on Snellen eye chart, or if they did not have normal color vision as judged by the Ishihara test (Clark, 1924).

### Visual stimuli and task

Visual stimuli were created using the Metropsis system (Cambridge Research Systems). This commercial apparatus for psychophysics implements tests in the Psykinematix language and uses a high-bit depth display with a 120 Hz refresh rate. Photometric calibration of the display was performed with a PR670 spectroradiometer (Photo Research). The stimulus was a circular field, 50-degrees of visual angle in diameter, with the outer edge smoothed by a Gaussian envelope (5 degrees SD; Figure 1A). Outside of the stimulus field, the display was set to the half-on primaries.

**Figure 1:**
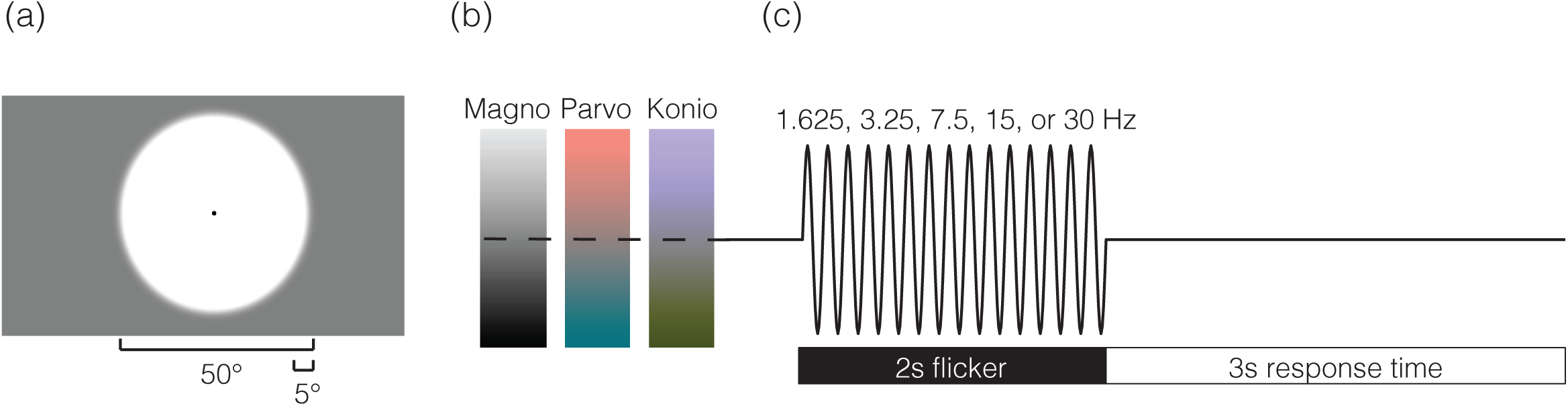
Spatial, temporal and spectral properties of visual stimuli. (a) The spatial structure of the stimulus consisted of a 50-degree diameter circle, with a 5-degree Gaussian envelope applied to the edge. The entire screen, and the stimulus background, was set to a midpoint gray. A 0.2-degree black circle was located in the center of the screen to aid fixation and obscure the foveal blue scotoma (Magnussen, Spillmann, Stürzel, & Werner, 2001). (b) Stimuli consisted of three spectral modulations that targeted the magnocellular (“black-white”), parvocellular (“red-green”), and koniocellular (“blue-yellow”) pathways. (c) Stimuli flickered sinusoidally at a rate of 1.625, 3.25, 7.5, 15, or 30 Hz. Two-second periods of flicker were followed by a 3-second response window during which the midpoint gray screen returned.

The spectral content of the stimulus field was modulated in time following a sinusoidal profile to create time-varying contrast that targeted the magno, parvo, and koniocellular pathways (Figure 1B; also see Supplementary Figure 1 for predicted and measured spectra). We used the method of silent substitution to target cone classes alone or in combination. Our estimates of photoreceptor spectral sensitivities were as previously described (Spitschan, Datta, Stern, Brainard, & Aguirre, 2016), and accounted for field size and age (Comission Internationale de L’Eclairage (CIE), 2005). The estimates assumed a 32 year-old observer with a 2mm diameter pupil. Modulation spectra were defined around the half-on (59 cd/m2) primaries and designed to produce isolated contrast on the LMS, L–M, and S mechanisms, and thus preferentially stimulate the magno, parvo, and koniocellular pathways, respectively.

Predicted cone spectral sensitivity varies as a function of eccentric field position due to the effect of macular pigment. Failure to account for this effect would produce differential contrast upon the targeted cone mechanism as a function of eccentricity, and inadvertent contrast upon nominally silenced mechanisms. To account for this, we varied the chromatic spectral modulations on the screen as a function of eccentricity. Although the CIE standard specifies fundamentals only for field sizes up to 10 degrees, we obtained the 30-degree estimates by extrapolation as described previously (Spitschan, Aguirre, & Brainard, 2015). The result was nominal Michelson contrast of 90%, 5.9%, and 81% upon the magnocellular, parvocellular, and koniocellular channels, respectively, that was spatially uniform across the stimulus field.

These contrast levels represent 90% of the maximum possible on the display given its gamut. The 10% “headroom” was reserved to allow for stimulus adjustment by flicker photometry to null residual luminance in the chromatic modulations for each subject. Subjects were shown the 50-degree sinusoidal flickering field that modulated around the background gray at 30 Hz. Subjects were asked to adjust the stimulus field to null residual luminance flicker by pressing the ‘up’ or ‘down’ arrow on the keypad, which added or subtracted 0.05 from the R, G, and B primary values. This test was repeated twice, once from below the estimated target value, and once from above. The value from each test was averaged to determine the nulling correction for the parvocellular and koniocellular channels. There was no difference between nulling values in the MwA and HAf groups (Supplementary Figure 2). One subject (MELA_0201) had difficulty following the nulling procedure. For this subject, the median nulling values across subjects was used instead.

On each of many trials, the half-on background was replaced with the stimulus field which flickered for 2 seconds at one of 5, log-spaced temporal frequencies (1.625, 3.25, 7.5, 15, and 30Hz; Figure 1C). This was followed by a 3-second response window during which the half-on background was again presented. Subjects were asked to rate the visual discomfort produced by the flicker on a 0 to 10 scale, with 0 being not uncomfortable at all, 5 being moderately uncomfortable, and 10 being extremely uncomfortable. The verbal response of the subject was recorded with a microphone and subsequently transcribed by the experimenters. A given block of 35 trials targeted a particular post-receptoral mechanism (magno, parvo, or konio), and the frequency order was pseudo-randomized within a block. Each block was run three times for a total of 21 repeats per stimulus condition.

Following our pre-registered protocol, we omit from the primary figures the results for the 15 Hz koniocellular stimulus. This stimulus produced an unexpected, spatially structured “brightness” modulation. We attempted to determine the source of this percept, but were ultimately unsuccessful. This observation could not be accounted for by luminance artifact from the monitor (Supplementary Figure 3). We consider it possible that the effect arises from the spatial variation that we introduced into the stimulus to account for macular pigment, interacting with longitudinal chromatic aberration (Taveras Cruz, He, & Eskew, 2019). As we were not confident in the properties of this particular stimulus, we elected to collect data for the modulation, but not include the data in tests of our hypotheses. The omitted data for the 15 Hz koniocellular stimulus are shown in the supplementary material (Supplementary Figure 4). Prior to beginning the experiment, ambient room light was adjusted to achieve a pupil size of ∼2.5 mm. Subjects were positioned in a chin rest 400 mm away from the screen.

### ssVEP recording

Prior work comparing fMRI and VEP suggests that ssVEPs are generated predominantly from primary visual cortex and area MT (Di Russo et al., 2007). We recorded ssVEP with a single active electrode over Oz based on the 10-20 international criteria for EEG placement over primary visual cortex. A ground and reference electrode were placed on each mastoid. The ssVEP signal was recorded using a biopac ERS100c amplifier with a maximum bandwidth of 1 Hz-10K Hz of at a 2K Hz sampling rate.

### Data analysis

Data were analyzed using publicly available (https://github.com/gkaguirrelab/vepMELAanalysis), custom MATLAB software. The signal was subject to a 0.5 to 150 Hz bandpass filter to remove non-physiologic oscillations, and a bandstop filter at 60 Hz to remove electrical line noise. Trials with any time point >0.08mV, or all values <0.02mV were removed to eliminate noisy trials due to poor electrode placement; this constituted 152 out of a total of 6,300 trials in the study (2.4%). The median response across trials in the time domain provided the cortical VEP. The first 0.5 seconds of the stimulus presentation were discarded to eliminate the onset response, leaving the remaining 1.5-second epoch. The signal was converted from the time domain into the frequency domain using a discrete Fourier transform. Responses contained a peak at the fundamental flicker frequency of the visual stimulus, which can be seen in the power spectral density (PSD) plots (supplementary figure 5). Prominent higher harmonic responses are also evident. Each PSD was fit using a previously described technique (Haller et al., 2018) to estimate and remove the aperiodic (non-oscillatory) component of the ssVEP (Supplementary Figure 5). Consistent with previous reports (Haller et al., 2018), the aperiodic signal was greatest at low frequencies and is well described by 1/frequency function. There were no significant differences in the aperiodic signal between groups or stimulus conditions (Supplementary Figure 6). The aperiodic fit was subtracted from the original signal to obtain the periodic signal (Supplementary Figure 5). Median responses for the fundamental flicker frequency of each stimulus were calculated across groups. A two-sample t-test was used for subject demographic comparison. The median responses for each group and spectral direction were used to fit a difference-of-exponents model that describes the temporal sensitivity (Hawken, Shapley, & Grosof, 1996). Temporal sensitivity fits were used to calculate the peak frequency and peak amplitude for each curve. Estimates of the variability of these measures in our population were obtained by repeating the fitting over 1000 bootstrap resamplings (with replacement) and calculating 95% confidence intervals.

In supplemental analyses we examined the response at both the fundamental and higher harmonic frequencies. The sum of the fundamental frequency and the 2F harmonic was calculated. As the 2F harmonic of the 30 Hz stimulus overlaps with powerline noise at 60 Hz, it could not be measured directly. Instead, the amplitude of the harmonic response was estimated. We observed that there was a linear relationship between the frequencies of the 1^st^, 2^nd^, 3^rd^, and 4^th^ harmonics, and the response amplitudes. Thus, the amplitude of response at the 30, 90, and 120 Hz frequencies was used to estimate the response at 60 Hz in this analysis.

In supplemental analyses we also tested for the presence of narrowband gamma oscillations. This was done by calculating Thomson’s multi-taper PSD estimate for each trial and taking the median value across trials.

## Results

We collected discomfort ratings and ssVEP data from 20 participants while they viewed high contrast, uniform, wide field flicker of varying temporal frequency that targeted the three different post-receptoral pathways (Figure 1). Below, we describe spectral modulations that target the magno, parvo, or koniocellular pathways as having a particular post-receptoral “direction”.

We derived the median discomfort rating across participants for each stimulus direction as a function of flicker frequency and derived peak frequency and peak amplitude from fitting a difference-of-exponentials model to the data (Hawken et al., 1996). There were no significant differences between MwA and headache free groups for these metrics for any of the post-receptoral directions directions (Supplementary Table 1). Therefore, we combined the data across the two groups. Supplementary Figure 6 provides the results separated by group.

### Discomfort sensitivity to flicker varies by post-receptoral pathway

Visual discomfort ratings varied by flicker frequency and stimulus direction (Figure 2a). Flicker targeting the magnocellular pathway was found to be most uncomfortable for the highest frequencies presented as compared to stimulation targeting the parvo and koniocellular pathways. The estimated temporal peak (and 95% confidence interval) of discomfort sensitivity was 18.6 Hz [16.5-19.4 Hz] for the magno, 10.3 Hz [9.5-11.6 Hz] for the parvo, and 9.1 Hz [7.8-10.7 Hz] for the koniocellular pathway (Table 2). Peak discomfort ratings also varied overall for the different stimulus directions. The greatest degree of discomfort was evoked by flicker directed at the magnocellular pathway: the median (across subject) peak amplitude of discomfort for the magnocellular pathway was 6.3 [5.5-8.0] compared to 4.0 [4.0-5.5] for the parvo and 4.5 [3.0-5.0] for the koniocellular pathway.

**Figure 2:**
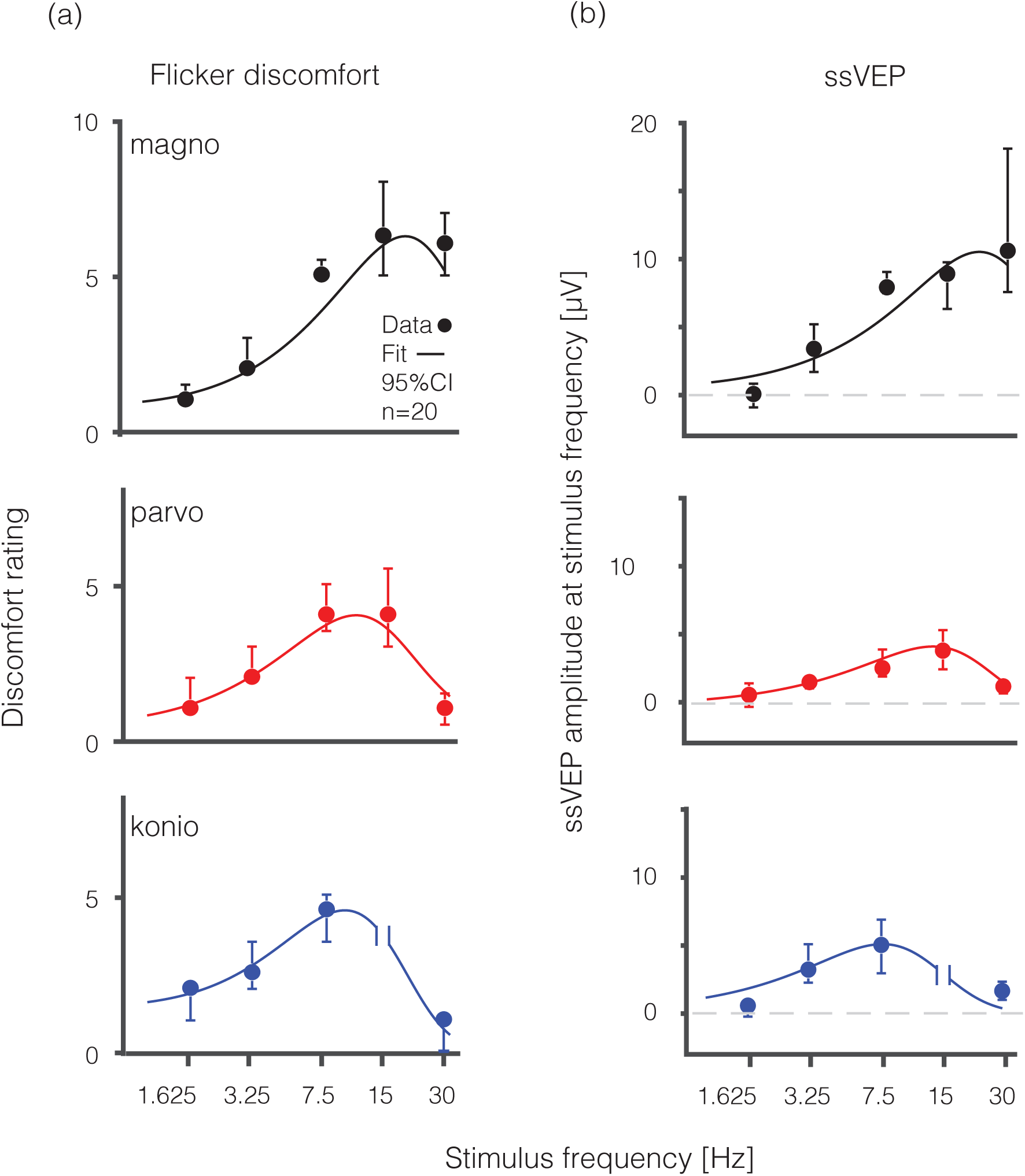
Visual discomfort ratings and visual cortex evoked responses across temporal frequency and spectral modulations. Median visual discomfort ratings on a 0-10 scale (a) and visual evoked response at the fundamental stimulus frequency represented in mV (b) are shown as a function of temporal frequency (Hz) for magnocellular (black), parvocellular (red), and koniocellular (blue) flickering stimuli. Measurements from the koniocellular stimulus flickering at 15 Hz were omitted (following our pre-registered protocol) as this stimulus was accompanied by a prominent, spatially structured “brightness” percept that we were unable to remove. Data are collapsed across HAf (n=10) and MwA (n=10) subjects. Error bars represent 95% confidence interval by bootstrap analysis. Fit line is derived from a difference-of-exponentials function.

**Table 2:** Peak frequency aind peak amplidute from temporal sensitivity difference of exponentials fit with median value and 95% confience interval by bootstrap analysis with replacement for the three spectral directions (magno, parvo, and koniocellular).

| | Peak Frequency [Hz]<br>Median [95% CI] | Peak Amplitude [μV]<br>Median [95% CI] |
| --- | --- | --- |
| Flicker<br>discomfort | Magno 18.6 Hz [16.5-19.4]<br>Parvo 10.3 Hz [9.5-11.6]<br>Konio 9.1 Hz [7.8-10.7] | Magno 6.3 [5.5-8.0]<br>Parvo 4.0 [4.0-5.5]<br>Konio 4.5 [3.0-5.0] |
| ssVEP | Magno 21.8 Hz [16.5-37.9]<br>Parvo 13.0 Hz [11.6-15.3]<br>Konio 9.1 Hz [5.7-9.5] | Magno 10.0 μV [7.4-18.0]<br>Parvo 4.1 μV [2.7-5.3]<br>Konio 4.9 μV [3.4-6.7] |

### Visual cortex response to flicker varies by post-receptoral pathway

The dependence of the ssVEP upon temporal frequency and stimulus direction was very similar to the relationship seen for visual discomfort (Figure 2b). Flicker directed at the magnocellular pathway evoked the strongest response for the highest frequencies presented, while peak response occurred at successively lower frequencies for the parvo and koniocellular pathways, respectively. The estimated peak of the visual evoked response (and 95% confidence intervals) was 21.8 Hz [16.5-37.9 Hz] for the magno, 13.0 Hz [11.6-15.3] for the parvo, and 9.1 Hz [5.7-9.5] for the koniocellular pathway, all similar to the flicker discomfort data (Table 2). This variation in peak neural response across temporal frequency is similar to what has been observed using functional magnetic resonance imaging (Spitschan et al., 2016). The amplitude of cortical response also varied for the different stimulus directions. The largest cortical response at 10 μV [7.4 - 18] was evoked by flicker targeting the magnocellular pathway, as compared to 4.1 μV [2.7-5.3] for the parvocellular pathway and 4.9 μV [3.4-6.7] for the koniocellular pathway.

### Flicker discomfort correlated with neural response in the primary visual cortex

As the evoked visual cortical response increased, so too did the magnitude of reported discomfort (Figure 3). Neural responses at the fundamental stimulus frequency were highly correlated with the level of reported discomfort from the flicker across the 14 stimulus conditions (R^2^ = 0.85, p=1.2e^-4^). This relationship did not appear to be unique to a single post-receptoral channel. The sum of the fundamental and harmonic responses showed a similar relationship (Supplementary Figure 7).

**Figure 3:**
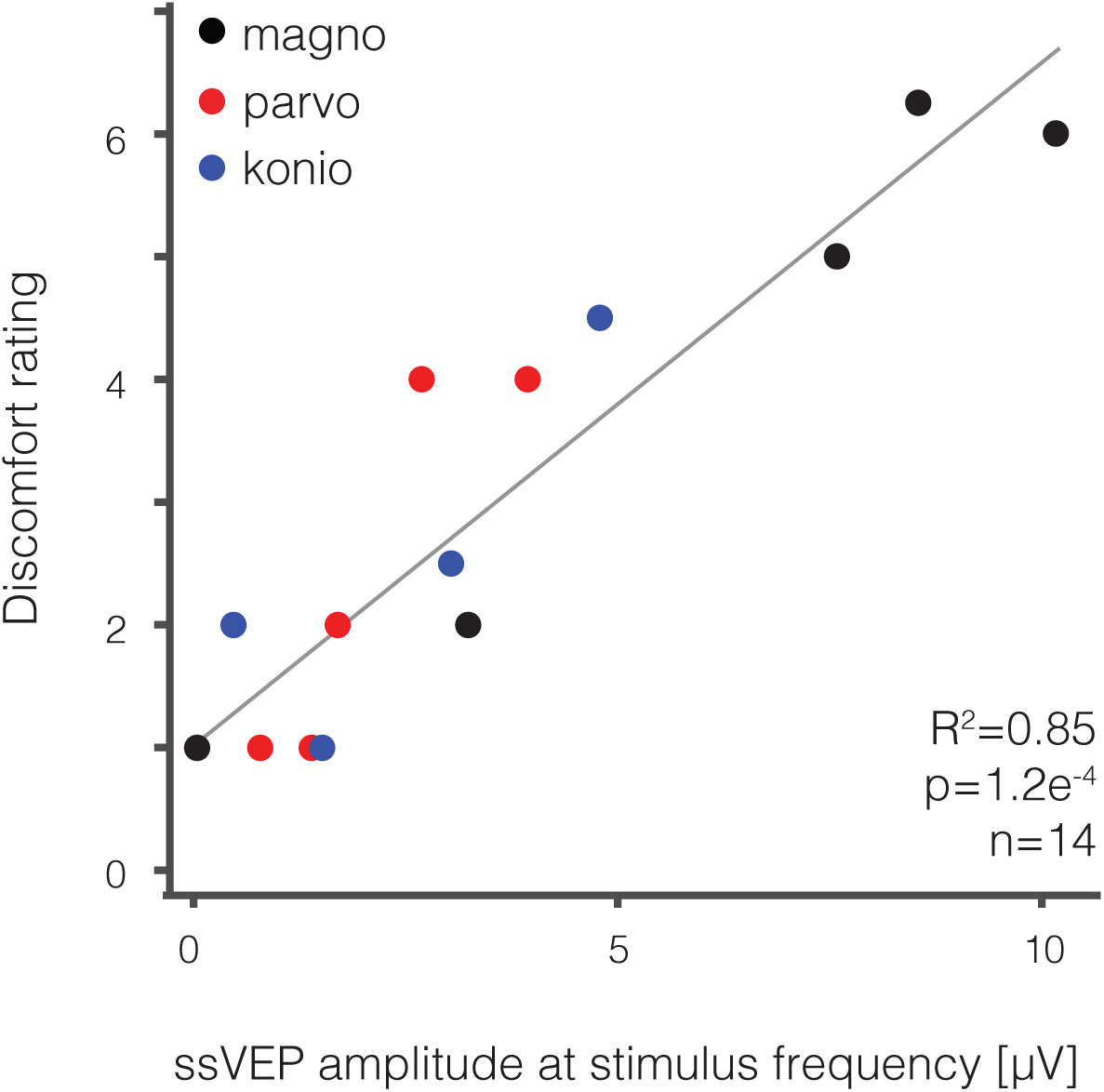
Visual discomfort strongly correlates with visual evoked response. The median visual discomfort rating (0-10 scale) is plotted as a function of the median visually evoked response (in μV) for each of the 14 unique stimuli designed to stimulate the magnocellular (black), parvocellular (red), and koniocellular (blue) pathways. There is a strong correlation between visual discomfort and visually evoked response (R^2^ = 0.85, p=3e^-6^).

## Discussion

### Summary

Our study demonstrates that visual discomfort elicited by flickering light correlates with evoked responses in primary visual cortex. This measurement supports a generally accepted theory that until now has seen limited, direct empirical support: that visual discomfort is the product of large amplitude neural responses within visual cortex (Aurora & Wilkinson, 2007). Our data also demonstrate that the relationship between cortical response and visual discomfort is independent of a particular post-receptoral pathway. Our findings are consistent with prior work that has examined variation in the detectability and salience of stimulus contrast directed at the post-receptoral pathways (Switkes, 2008). While there have been reports that different colors of light evoke greater discomfort (Noseda et al., 2016), it is important to note that cone signals are only experienced by the central nervous system through the coding of the retinal ganglion cells and post-receptoral channels. We can say that visual discomfort from flicker is not the domain of any particular ‘color’ or chromatic contrast for the examined stimulus regime. This finding suggests that the relationship between cortical activity and discomfort is not limited to a particular post-receptoral pathway, but rather reflects a more general phenomenon of the visual system.

### Comparison with prior studies

While several studies have examined the effect of stimulus variation upon visual cortex neural response or upon reports of visual discomfort, few studies have combined these measurements. Haigh and colleagues (2013) demonstrated that parametric variation in the color separation of isoluminant gratings was related both to greater reported discomfort and greater visual cortex hemodynamic responses (as measured by near-infrared spectroscopy). In our work we tailored our stimuli to specifically target the post-receptoral channels and examined the effect of variation in flicker frequency with nominally constant stimulus contrast.

Prior studies that have made use of ssVEP have found a weak (O’Hare, 2017) or negative (Bjørk, Hagen, Stovner, & Sand, 2011) correlation between visual discomfort and visual cortical response. We believe that an important difference between this prior work and our study is the isolation of the induced periodic signal component from the aperiodic component (Haller et al., 2018). Beyond reducing noise, accounting for the aperiodic signal prevents low temporal frequency stimuli from appearing as if they induce larger cortical responses.

### Comparison of migraine with aura and headache free participants

There was minimal difference in visual discomfort ratings between the groups, and correspondingly, we did not find significant differences in cortical response between MwA subjects and HAf controls. This absence of a difference in cortical response in our study is contrary to prior studies that have found increased visual responsivity in migraine, particularly migraine with visual aura (Áfra, Cecchini, De Pasqua, Albert, & Schoenen, 1998; Aurora, Ahmad, Welch, Bhardhwaj, & Ramadan, 1998; Aurora, Barrodale, Chronicle, & Mulleners, 2005; Boulloche et al., 2010; Chen, Lin, et al., 2011; Chen, Wang, et al., 2011; Datta et al., 2013; Denuelle et al., 2011).

We might consider several possible explanations for this discrepancy. It may be the case that our MwA population has a relatively low disease burden, and thus a relatively normal cortical phenotype. Based on headache frequency in the prior three months, our MwA subjects have episodic migraine, not chronic, based on International Classification of Headache Disorders 3^rd^ Edition(International Headache Society, 2018). Our subjects did not use preventative migraine medications. Prior work suggests that increased cortical response accompanies increased disease burden (Chen, Wang, et al., 2011). While this makes it possible that our migraine subjects had too low a disease burden to demonstrate cortical hyper-responsiveness, we note that the headache burden and reported photophobia symptoms in our patient population was nonetheless equal or greater than that examined in some of these prior studies (e.g., Datta et al., 2013).

We think a more likely explanation is that differences between the migraine and control groups may not be present for the stimulus regime we examined. Our stimulus was spatially uniform, and had a maximum flicker frequency of 30 Hz. Prior studies that have found enhanced cortical responses in migraine have generally used stimuli with high-contrast spatial structure. It may be the case that the addition of spatial structure or more rapid flicker would have revealed a difference in cortical response between the groups. The magnocellular pathway is a particular candidate in which this difference could emerge. Our measurements track the parvocellular and koniocellular pathways past their peak response. Therefore, it is unlikely that a greater neural response would be present for flicker of higher temporal frequency directed at these chromatic channels. In contrast, the response to the magnocellular stimulus continued to rise at our highest temporal frequency. It is possible that an even greater neural response, and a potential difference between MwA and HAf subjects, might be found at higher temporal frequencies. In our data, cortical responses in the headache-free group to magnocellular stimulation appear to peak between 15 and 30 Hz, whereas responses in the MwA group continue to trend upwards, consistent with this theory (see Supplementary Figure 6A).

### Narrowband gamma oscillations

Narrowband gamma oscillations have been postulated as a neural correlate for visual discomfort. These resonant neural signals between 30-80 Hz (Buzsáki & Wang, 2012) are evoked by a restricted set of high-contrast stimuli (Hermes, Miller, Wandell, & Winawer, 2015; Dora Hermes, Miller, Wandell, & Winawer, 2015), and do not show strong correlation to multi-unit neuronal activity (Ray, Crone, Niebur, Franaszczuk, & Hsiao, 2008; Ray & Maunsell, 2011; Winawer et al., 2013). The visual features that evoke narrowband gamma are also those that tend to cause discomfort (Adjamian et al., 2004; Hermes et al., 2015; Hermes, Trenité, & Winawer, 2017). Narrowband gamma oscillations are enhanced in migraine (Coppola et al., 2007) suggesting they represent network dynamics involved in hypersensitivity to visual stimuli. While these signals can be measured in surface EEG data (Long, Burke, & Kahana, 2014), we did not find in our data evidence that our stimuli evoke narrowband gamma (Supplementary Figure 9).

### Pain pathways involved in photophobia

Our data do not assign a causal relationship between cortical activity and the sensation of discomfort. Multiple neural pathways have been implicated in visual discomfort, some of which involve the visual cortex, while others do not (for review, see Digre & Brennan, 2012).

Projections of melanopsin-containing intrinsically photosensitive retinal ganglion cells (ipRGCs) to the thalamus have been offered as a mechanism of light induced discomfort that does not involve the visual cortex (Noseda et al., 2010). However, this mechanism seems ill suited to explain discomfort from flicker, as the stimulus variation is quite rapid relative to the slow kinetic responses of the ipRGCs (Do et al., 2009).

### An “out of gamut” error

The widely accepted theory of “visual stress” posits that the experience of visual discomfort corresponds to overactivation of the visual cortex (Wilkins, 1995). Our work supports this idea, and further demonstrates that the relationship between cortical response and visual discomfort is general, operating across the three post-receptoral visual pathways. While consistent with the empirical data, the theory of visual stress does not provide an explanation for *why* larger amplitude visual cortex responses are aversive. We are unaware of any evidence that stimuli such as ours are actually injurious to the nervous system, or produce demands with which the neurometabolic or neurovascular system cannot cope. Instead of viewing neural activity as a physical stressor upon the central nervous system, we believe that a better way of understanding visual discomfort is as a signal regarding information processing.

It is our view that visual discomfort is just what it feels like to be a neural system in an efficient signal processing range. This “out of gamut” account of visual discomfort does away with the need to identify a putative physical injury caused by visual stimulation.

This view is consistent with a growing body of literature that has found that stimuli are perceived as unpleasant when their spatial and temporal content departs from the statistics of natural environments (Juricevic et al., 2010; Penacchio & Wilkins, 2015; Yoshimoto et al., 2017). The visual system is remarkably adaptive across long and short time scales, and an important function of that adaptation is to allow neurons to encode sensory information using a range of representation that is well matched to the current statistics of the environment (Barlow, 2001). Our stimuli were distinctly unnatural (wide-field, high-contrast, sinusoidal flicker), and thus well suited to place neural coding at a representational disadvantage.

## Conclusions

We find a linear relationship between visual discomfort and visual cortex response, providing empirical support to the long-standing theory of visual stress. Visual discomfort from temporal flicker is not the unique domain of a particular chromatic or achromatic post-receptoral pathway. Visual discomfort and visual cortical response did not differ substantially between MwA and HAf subjects for the studied stimuli.

## Acknowledgements

Acknowledgments

None This work was supported by National Institutes of Health Grant R01 EY024681(to G.K.A. and D.H.B.), Core Grant for Vision Research P30 EY001583, and Department of Defense Grant W81XWH-15-1-0447 (to G.K.A.).

---

**Figure S1:**
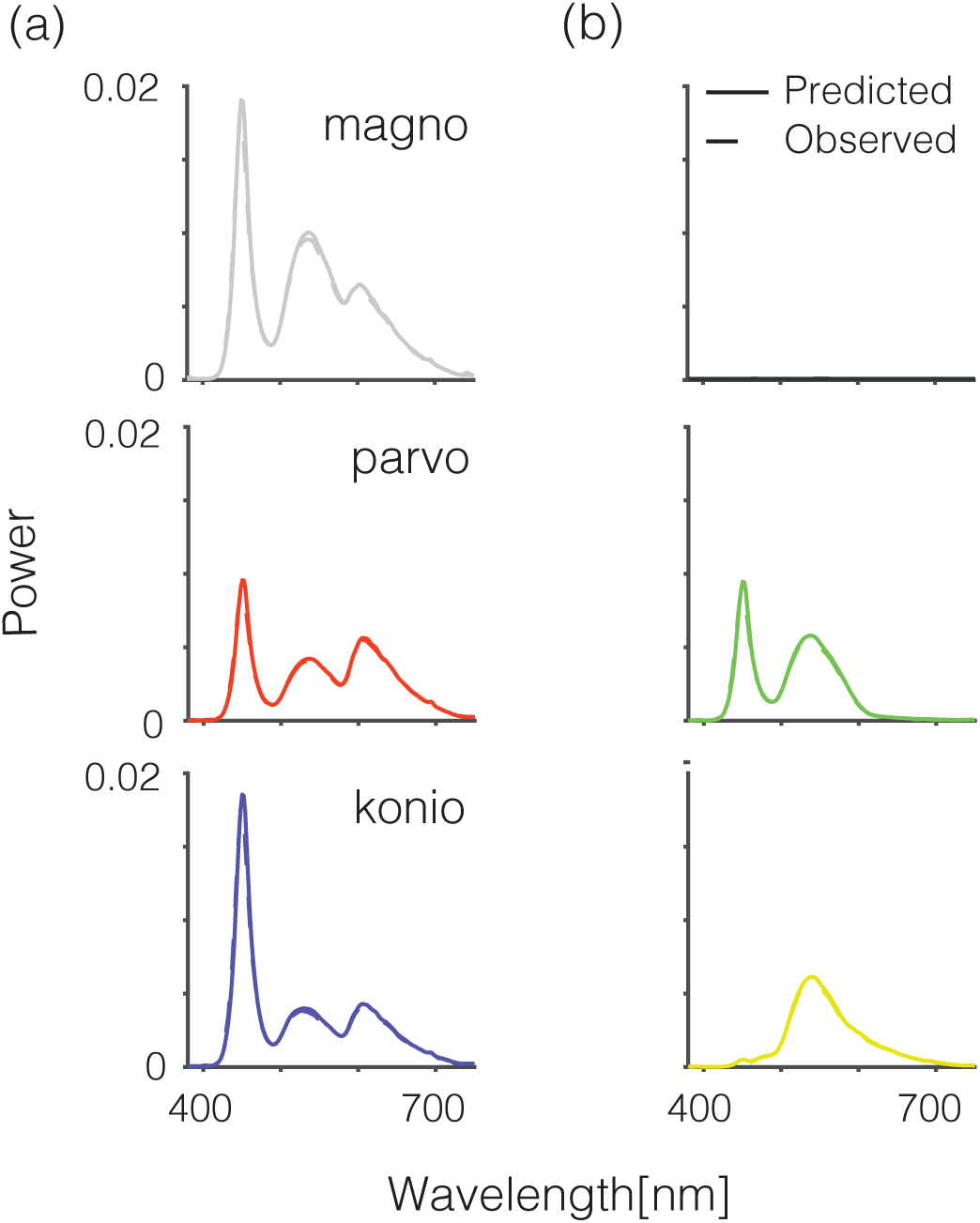
Power spectra for the nominal (solid line) and measured (dotted line) stimulus spectra for the positive (a) and negative (b) arms of the magnocellular (top), parvocellular (middle) and koniocellular (bottom) stimuli. The measured values are in close agreement with the predicted values.

**Figure S2:**
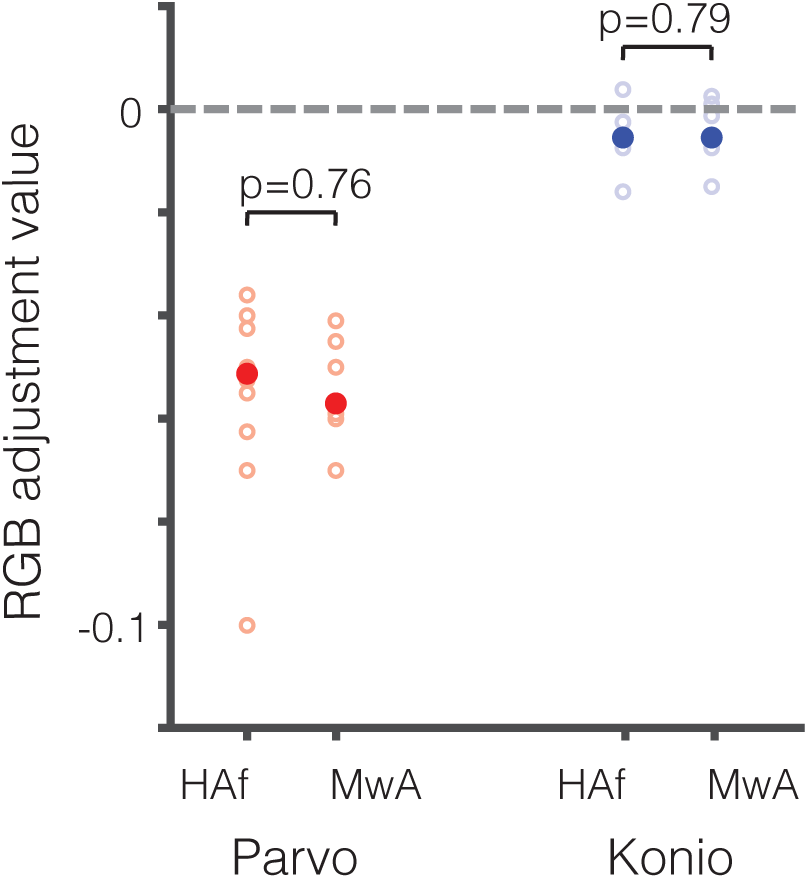
RGB nulling values for the parvocellular (red) and koniocellular (blue) stimuli for each subject (open circles). Median values for the HAf and MwA groups are shown (closed circles). P-values represent 2-way t-test.

**Figure S3:**
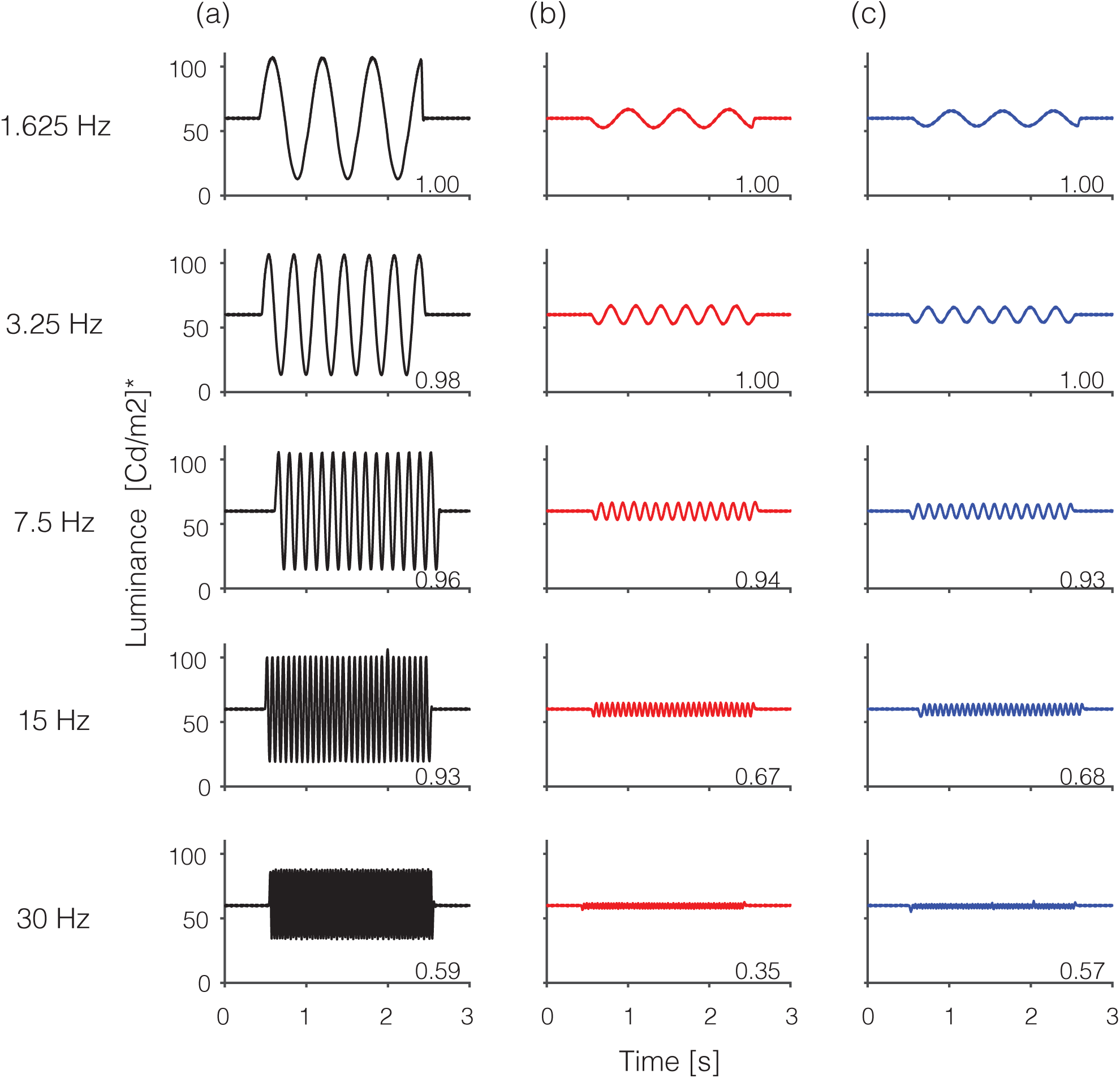
We measured the flickering stimuli in the center of our display using a photometer with a high temporal refresh rate (Klein 650, Klein Instruments Corporation). The photometer reports calculated “luminance” values in units of cd/m^2^. We measured the photometer-reported luminance of the stimuli across a 2-second period of flicker for the magno (a), parvo (b), and koniocellular (c) modulation directions across all temporal frequencies. It should be noted that the photopic luminosity function assumed by the device differs from the spectral sensitivity functions that we assumed for the purposes of generating our stimuli. As aconsequence, our nominally iso-luminant stimuli (those that target the parvo and koniocellular pathways) are reported by the Klein device to have a small temporal modulation of “Klein luminance”. We were able to use this modulation to confirm that 1) the stimuli retain their general sinusoidal form across temporal frequencies and modulation directions, however 2) there is a drop in the modulation amplitude with increasing stimulus frequency (proportional amplitude relative to the1.625 Hz stimulus is indicated). This later effect is comparable in size across the modulation directions, and likely represents the imperfect recreation of a high-frequency sinusoid at the 120 Hz refresh rate available for the monitor.

**Figure S4:**
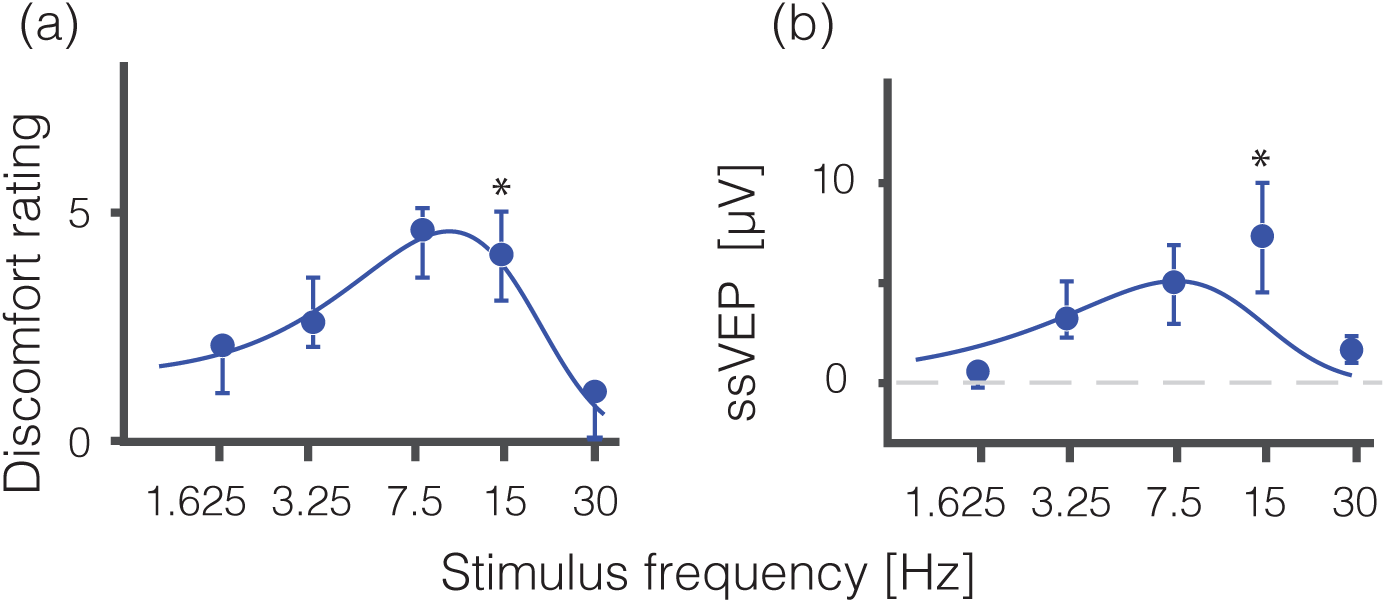
Data and fits from figure 2 plotted with the omitted visual discomfort and ssVEP values of the 15 Hz koniocellular stimulus.

**Figure S5:**
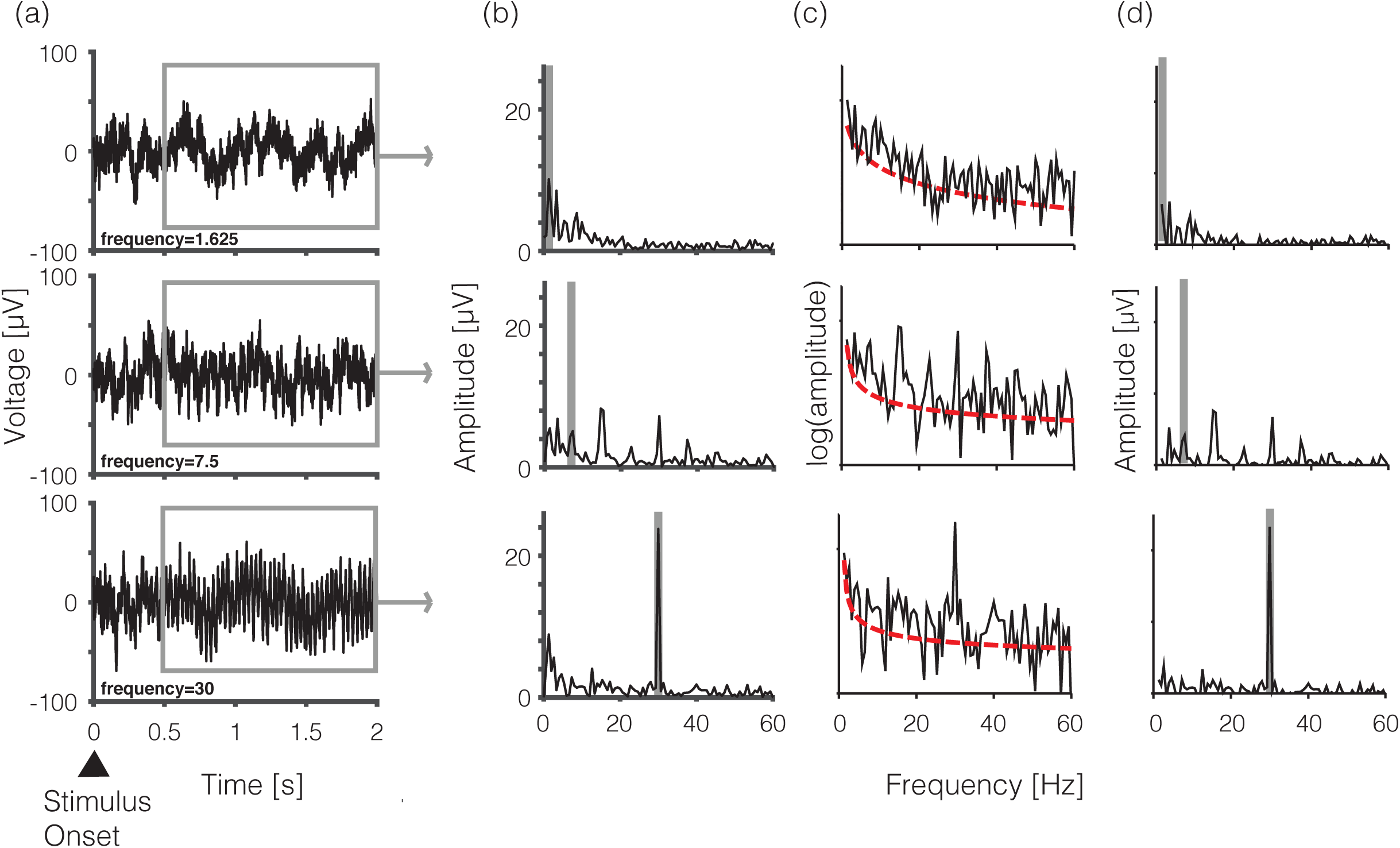
ssVEP analysis for an example subject. (A) median ssVEP response over the 2-second stimulus epoch for 1.625 Hz (top), 7.5 Hz (middle), and 30 Hz (bottom) magnocellular stimuli. Gray outline denotes the portion of the response epoch that was retained after discarding the response onset. (B) ssVEP signal converted from the time domain to frequency domain through discrete Fourier transform. Gray boxes indicate the fundamental frequency of the visual stimulus. (C) ssVEP signal amplitude in the frequency domain plotted on a logarithmic scale (black) with aperiodic component fit (red)(Haller et al., 2018). (D) The periodic component of the ssVEP signal in the frequency domain (i.e., with the aperiodic signal subtracted from the original signal).

**Figure S6:**
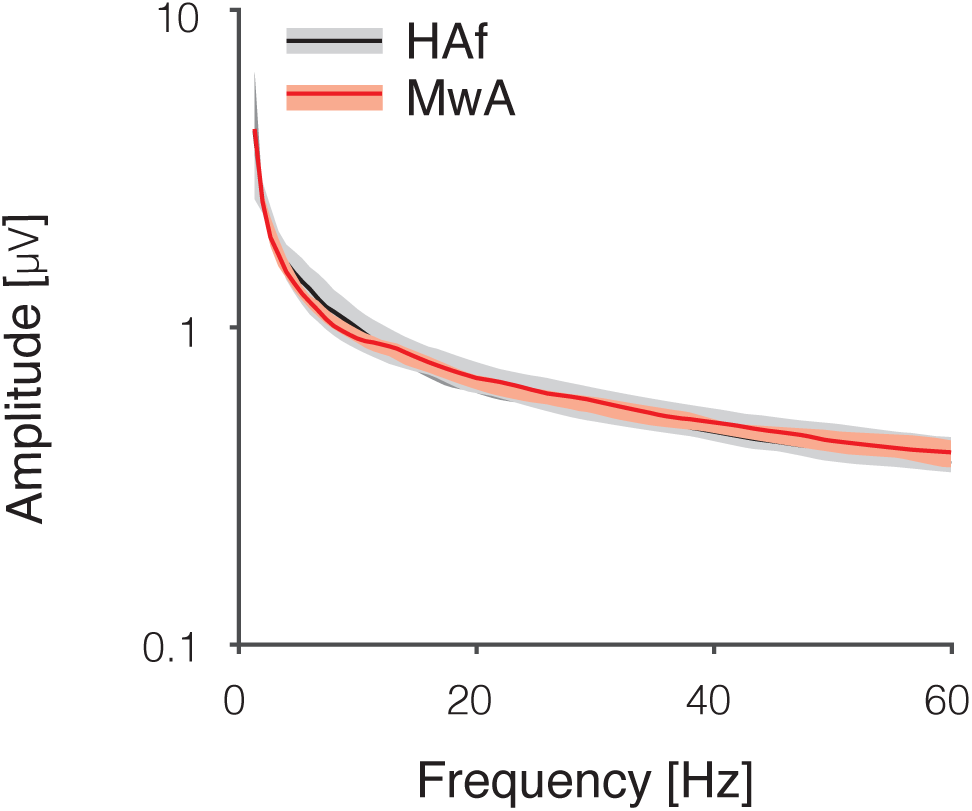
Median of the aperiodic fits across all stimuli for HAf (black) and MwA (red) subjects. Shaded area represents 95% confidence intervals.

**Figure S7:**
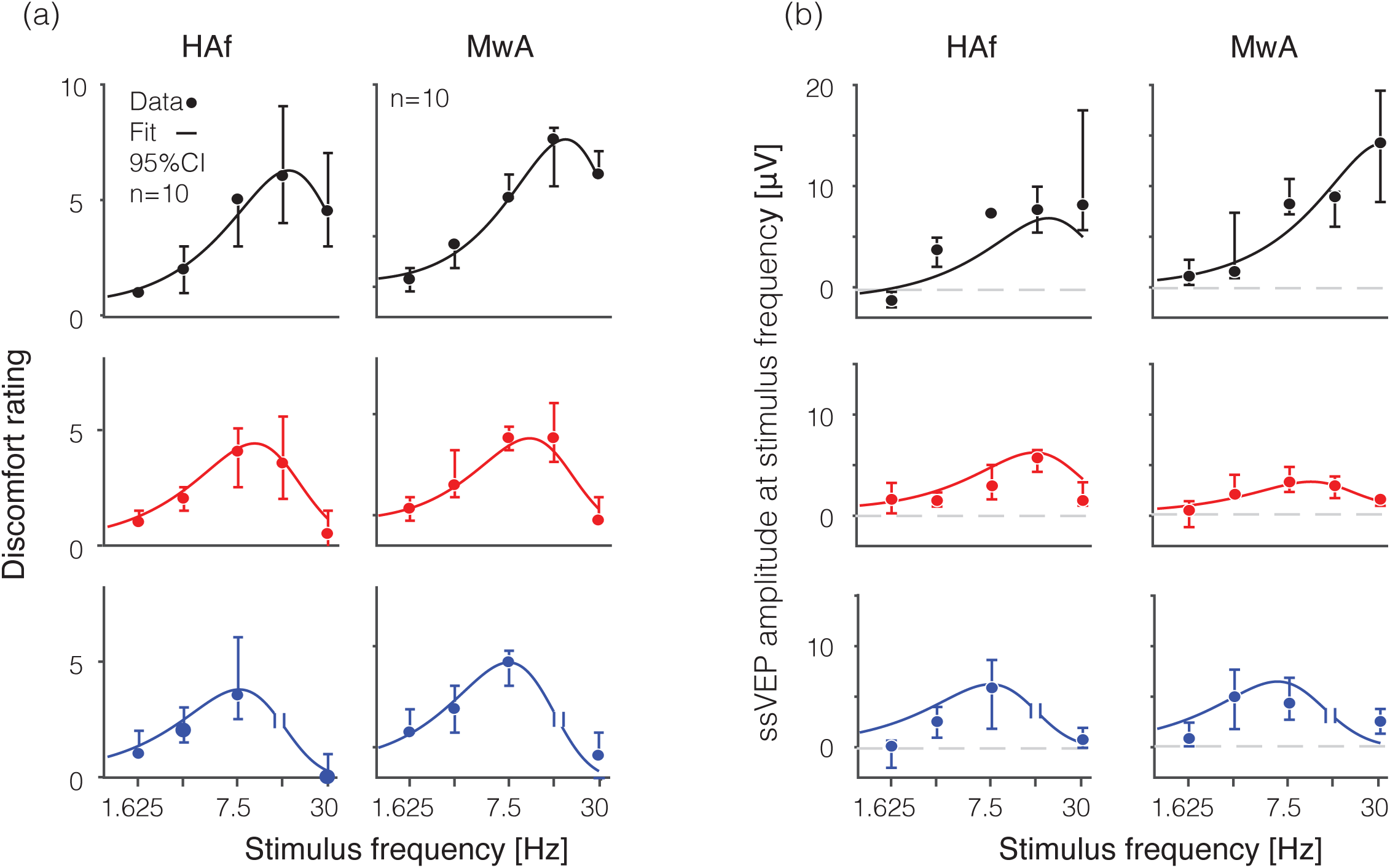
Visual discomfort ratings and visually evoked responses across temporal frequency and spectral modulations comparing HAf and MwA groups. Median visual discomfort ratings on a 0-10 scale (A) and visual evoked response at the fundamental stimulus frequency represented in mV (B) are shown as a function of temporal frequency (Hz) for magno (black), parvo (red), and koniocellular (blue) flickering stimuli. Measurements from the koniocellular stimulus flickering at 15 Hz were omitted (following our pre-registered protocol) as this stimulus was accompanied by a prominent, spatially structured luminance percept that we were unable to remove. Data are collapsed across HAf (n=10) and MwA (n=10) subjects. Error bars represent 95% confidence interval by bootstrap analysis. Fit line is derived from a difference-of-exponentials function.

**Figure S8:**
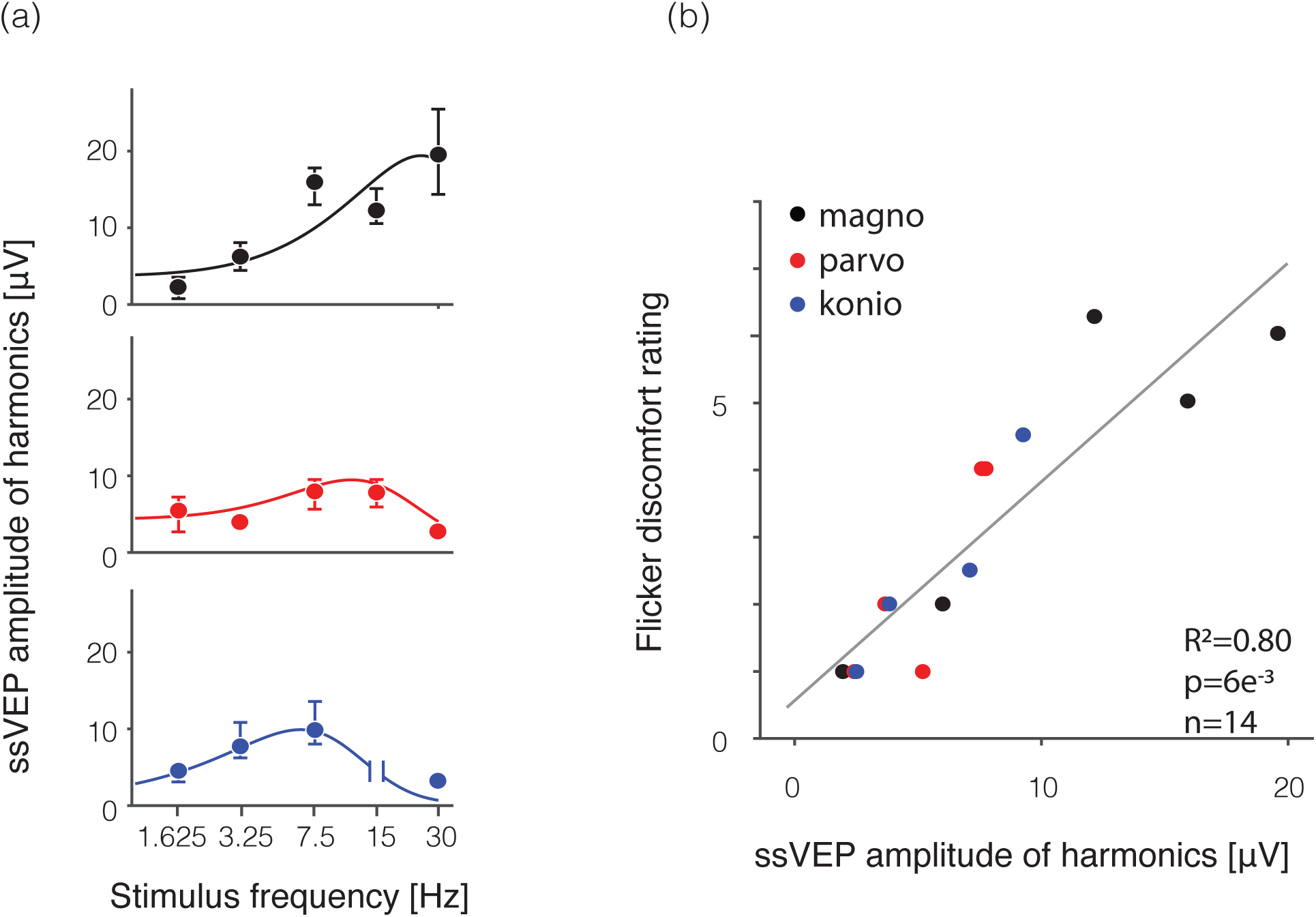
Stimulus frequency harmonics of the ssVEP signal. (A) shows the sum of the 1st and 2nd harmonics of the ssVEP signal for magnocellular (black), parvocellular (red), and koniocellular (blue) stimuli as a function of stimulus frequency. (B) shows the flicker discomfort rating as a function of the sum of fundamental and second harmonic ssVEP for the 14 stimuli.

**Figure S9:**
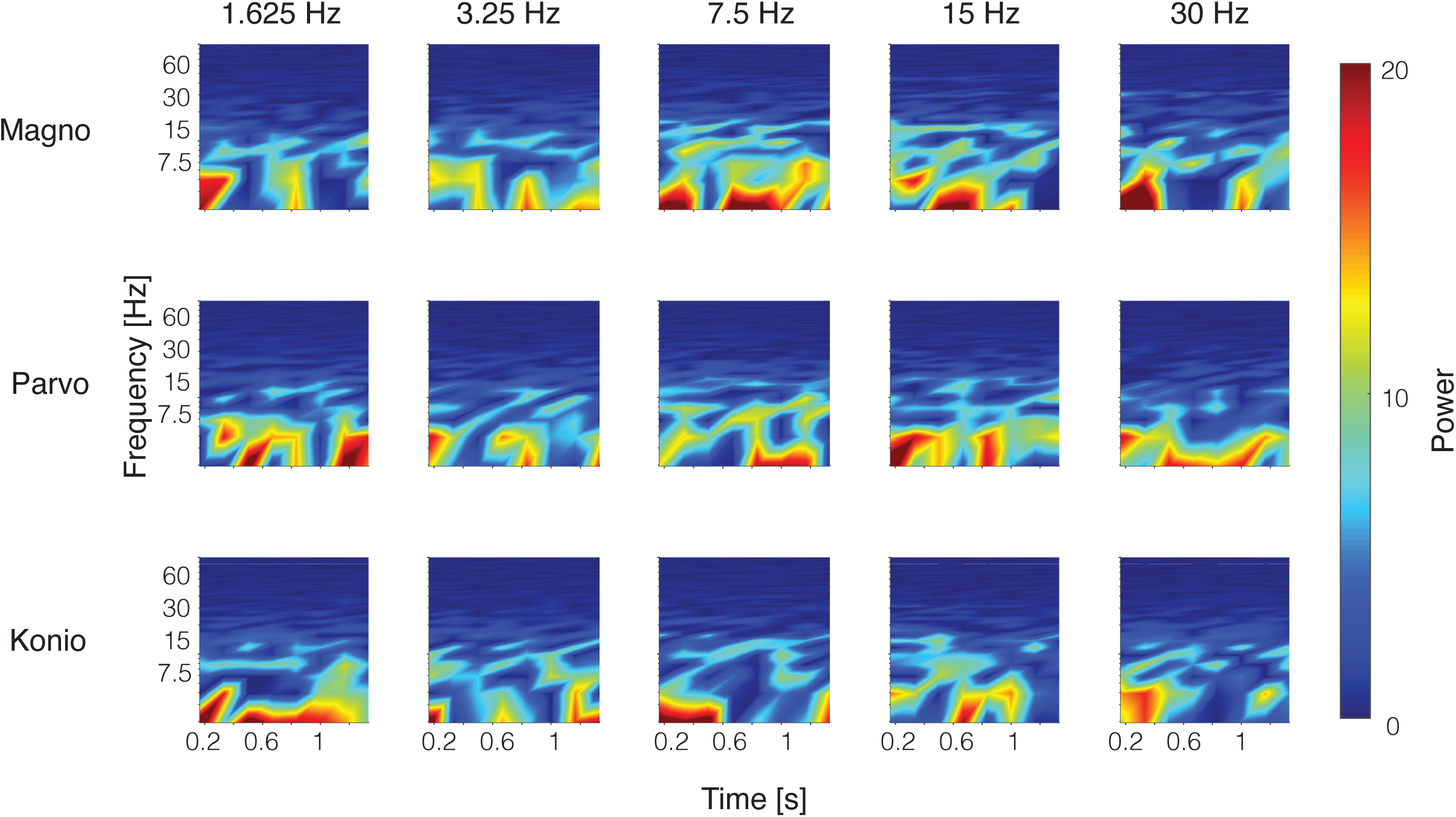
Narrowband gamma oscillations. Thomson’s multitaper power spectral density (PSD) calculated across all subjects for the magnocellular (top), parvocellular (middle), and koniocellular (bottom) stimuli across 5 temporal frequencies. PSD was calculated from 200ms after stimulus onset to 1500ms after stimulus onset to focus on the steady state signal.

**Table Si:** Peak frequency and peak amplidute from temporal transfer function difference of exponentials fit with median value and 95% confience interval by bootstrap analysis with replacement for HAf vs. MwA for the three spectral directions (magno, parvo, and koniocellular).

| | Peak Frequency [Hz]<br>Median [95% CI] | | Peak Amplitude [μV]<br>Median [95% CI] | |
| --- | --- | --- | --- | --- |
|  | MwA | HaF | MwA | HaF |
| Flicker discomfort | Magno 31.0 Hz [16.6-44.9]<br>Parvo 12.5 Hz [10.6-14.1]<br>Konio 9.3 Hz [7.6-14.1] | Magno 16.7 Hz [14.1-18.5]<br>Parvo 9.7 Hz [7.9-11.5]<br>Konio 8.5 Hz [7.6-9.7] | Magno 7.5 [6.0-8.0]<br>Parvo 4.5 [4.0-6.0]<br>Konio 5.0 [4.0-5.5] | Magno 6.0 [4.0-9.0]<br>Parvo 4.0 [2.0-5.0]<br>Konio 3.5 [2.5-5.0] |
| ssVEP | Magno 31.1 Hz [16.6-42.7]<br>Parvo 12.5 Hz [10.3-14.1]<br>Konio 6.4 Hz [5.5-9.5] | Magno 17.9 Hz [10.3-36.4]<br>Parvo 13.5 Hz [12.5-17.9]<br>Konio 9.5 Hz [5.9-9.5] | Magno 14.0 μV [8.9-19.0]<br>Parvo 3.2 μV [1.9-4.7]<br>Konio 4.9 μV [4.1-7.3] | Magno 7.6 μV [5.8-16.0]<br>Parvo 5.5 μV [4.2-6.3]<br>Konio 5.4 μV [1.5-8.4] |

